# Phenotypic Convergence in Bacterial Adaptive Evolution to Ethanol Stress

**DOI:** 10.1101/011577

**Authors:** Takaaki Horinouchi, Shingo Suzuki, Takashi Hirasawa, Naoaki Ono, Tetsuya Yomo, Hiroshi Shimizu, Chikara Furusawa

## Abstract

Bacterial cells have a remarkable ability to adapt and evolve to environmental changes, a phenomenon known as adaptive evolution. Adaptive evolution can be explained by phenotypic changes caused by genetic mutations, and by phenotypic plasticity that occur without genetic alteration, although far less is known about the contributions of the latter. In this study, we analyzed phenotypic and genotypic changes in *Escherichia coli* cells during adaptive evolution to ethanol stress. Phenotypic changes were quantified by transcriptome and metabolome analyses and found similar among independently evolved ethanol tolerant strains. The contribution of identified mutations in the tolerant strain was evaluated by using site-directed mutagenesis, which suggested that the fixation of these mutations cannot fully explain the observed ethanol tolerance. The phenotype of ethanol tolerance was stably maintained after an environmental change, suggesting that a mechanism of non-genetic memory contributed to at least part of the adaptation process.

## Introduction

Recent advances in high-throughput sequencing have made it possible to identify and study beneficial mutations in whole-genomic sequences during microbial adaptive evolution. For example, several mutations were identified as beneficial in adaptively evolved *E. coli* strains that used glycerol as the carbon source (Herring, et al. 2006). Other studies using laboratory evolution and genome resequencing have provided evidence that genomic mutations contribute to adaptive phenotypic changes against various environments, including several carbon sources (Conrad, et al. 2009; Cooper and Lenski 2010; Lee and Palsson 2010), different temperatures (Kishimoto, et al. 2010; Tenaillon, et al. 2012), high ethanol (Goodarzi, et al. 2010) and isobutanol (Atsumi, et al. 2010) stresses, and the presence of antibiotics (Toprak, et al. 2012). Furthermore, non-additive interactions between mutations, i.e. epistasis, significantly contribute to the dynamics of adaptive evolution (Khan, et al. 2011). The above studies argue that adaptive phenotypes arise through a process that involves natural selection and genotypic changes caused by mutations.

However, in addition, phenotypic plasticity, i.e., phenotypic changes without genetic alterations, can also contribute to environmental adaptation and evolution. A well known example of phenotypic plasticity in microorganisms is environmental stress response, in which the gene expressions are drastically changed in response to diverse environmental changes. Although phenotypic changes by such non-genetic mechanisms generally have much shorter time-scales than evolutionary dynamics, several works suggest that phenotypic plasticity strongly affects phenotypic changes caused by mutations. The earliest support was presented by Waddington (Waddington 1959, 1957). He showed that the artificial selection of a phenotype resulting from phenotypic plasticity can be easily stabilized by genetic mutations. Similar observations were demonstrated in other evolutionary dynamics (Eldar, et al. 2009; Rutherford and Lindquist 1998; Suzuki and Nijhout 2006), which suggests that phenotypic plasticity facilitates novel adaptive phenotypes caused by genetic mutations (Baldwin 1896; Kaneko and Furusawa 2006). Quantitative understanding of the phenomenon, however, is lacking. For this purpose, greater analysis is needed on genetic and non-genetic contributions to adaptive phenotypic changes during adaptive evolution.

In this study, we analyzed phenotypic and genotypic changes in the laboratory evolution of *E. coli* cells under ethanol stress. Using six independently evolved ethanol tolerant strains from our previous study (Horinouchi, et al. 2010), we quantified phenotypic changes by transcriptome and metabolome analyses to evaluate how adaptive phenotypic changes are similar among the different strains. Furthermore, we assessed genotypic changes in the tolerant strains using high-throughput sequencers. Finally, we introduced all the identified mutations in the genome of the parent strain into one of the tolerant strains and evaluated how these mutations contributed to adaptive phenotypic changes. The relationship between the timing of mutation fixation and phenotypic changes was also analyzed.

## Results

### Time-series expression analysis in adaptive evolution under ethanol

We previously obtained 6 independently evolved ethanol tolerant *E. coli* strains, strains A through F, by culturing cells under 5% ethanol stress for about 1000 generations and found a significantly larger growth rate than the parent strains (Horinouchi, et al. 2010). To elucidate the phenotypic changes that occurred during adaptive evolution, first we quantified the time-series of the expression changes by microarray analysis. Starting from frozen stocks obtained at 6 time points in laboratory evolution (arrows in Figure 1a), cells were cultured under 5% ethanol stress, and mRNA samples were obtained in the exponential growth phase for microarray analysis (complete data are presented in Table S1).

**Figure 1.**
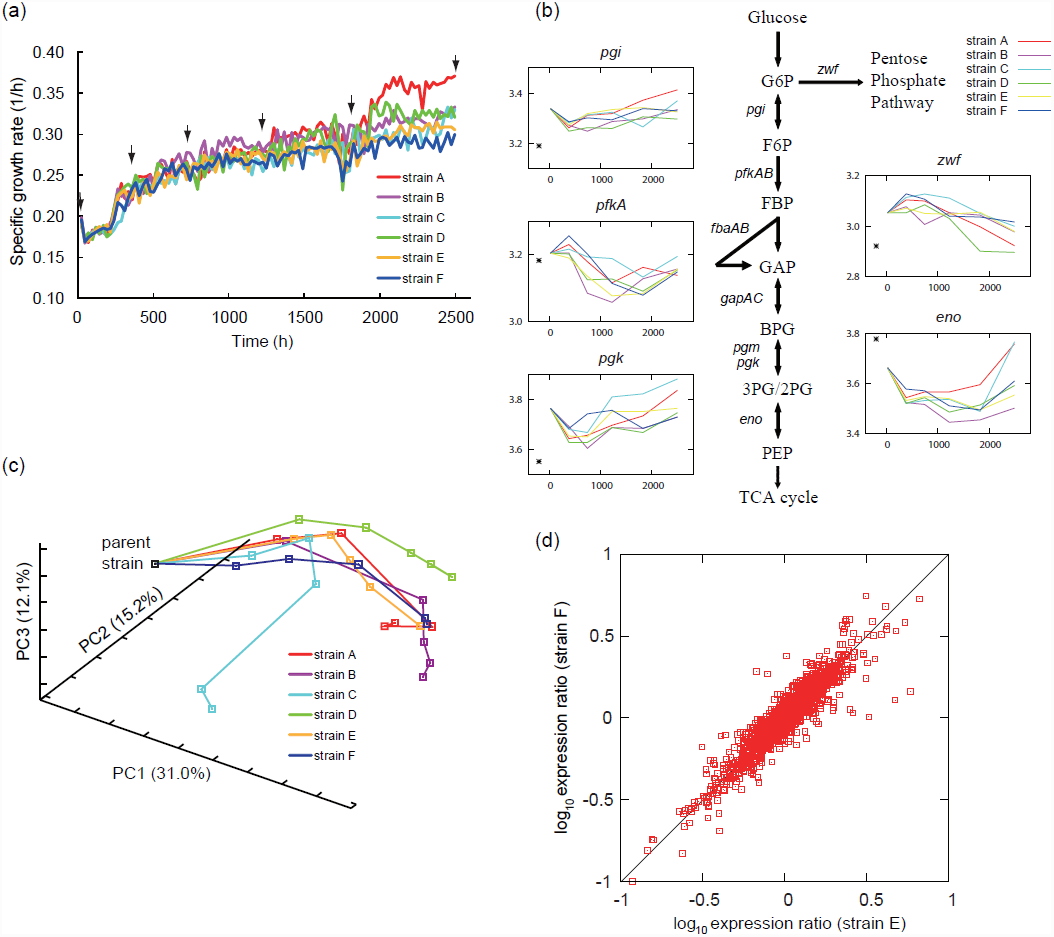
Time-series transcriptome analysis for adaptive evolution of *E. coli* to ethanol stress. (a) Time-course of the specific growth rate in evolution experiments. Arrows indicate the time-point at which mRNA samples were collected for transcriptome analysis (0, 384, 744, 1224, 1824, and 2496 hours after starting the culture). (b) Expression changes of 5 genes (*pgi, pfkA, pgk zwf*, and *eno*), all related to upper glycolysis. In each inset, the horizontal axis shows time (hours), and the vertical axis shows expression level (a.u.). Asterisks (*) indicate expression levels obtained absent ethanol stress as a reference. The expression changes of representative genes in the central metabolic pathway including glycolysis, the pentose phosphate pathway, and TCA cycle are presented in Figure S1. (c) Changes in PCA scores during adaptive evolution. Starting from the parent strain, changes in the expression profiles during adaptive evolution are plotted as orbits in the three-dimensional PCA plane. (d) Correlation between expression changes that occurred in strains E and F. Horizontal and vertical axes are log-transformed expression ratios with the parent strain, and each dot represents the expression change of the gene.

The results of the time-series transcriptome analysis revealed that the expression changes during adaptive evolution were similar among tolerant strains. As an example, Figure 1b shows the expression changes of genes in upper glycolysis (the expression changes of other genes in the central metabolic pathway are presented in Figure S1). Interestingly, common expression changes were not always monotonic (e.g., *pfkA* gene) over time, but were rather synchronized complex expression changes on a much longer time-scale than the generation time observed. Additionally, a common and gradual up-regulation of genes involved in some amino acid biosynthesis pathways was observed (Figure S2). Our previous work suggests these pathways might contribute to ethanol tolerance (Horinouchi, et al. 2010).

Figure 1c shows overall expression changes during the adaptive evolution of the six tolerant strains by principal component analysis (PCA). The orbits in PCA space, which represent expression profile changes, were similar except for strain C. The reason for this exception might be that strain C has an approximately 200k-bp region in the genome that was duplicated (discussed below), and the expression levels of genes in this region was increased by this duplication. The expression analysis also demonstrated that the overall expression changes between the parent and tolerant strains at the end-point were quite similar (Figure 1d and Figure S3). These results indicated that even though these strains adapted to ethanol stress in independent cultures, the expression profiles converged into almost identical adapted states with similar orbits of expression changes.

### Metabolome analysis of ethanol tolerant strains

To further characterize the phenotypic changes that occurred in the tolerant strains, we measured metabolite concentration changes between parent and tolerant strains. Using capillary electrophoresis time-of-flight mass spectrometry (CE-TOFMS), we quantified the intra-cellular concentrations of 83 metabolites (complete data are presented in Table S2). The intra-cellular concentrations of some amino acids in the parent and tolerant strains are presented in Figure 2a. These concentrations generally decreased in the tolerant strains, except for that of methionine. The decrease was especially true for amino acids that originated from precursors in the tricarboxylic acid (TCA) cycle. This fact might suggest a decrease of metabolic flux in the TCA cycle in tolerant strains, a conclusion supported by the significant decrease in the expression of genes related to the TCA cycle (Figure S1). For example, glutamate acts as a major amino-group donor in amino acid biosynthesis, and thus a decrease in its concentration can cause a decrease in the concentration of other amino acids. The decrease in amino acid concentration can activate the amino acid starvation response, which is consistent with the up-regulation of genes related to amino acid biosynthesis. In contrast, the concentrations of metabolites in purine metabolism generally increased (Figure S4). The concentration increase here might be caused by the up-regulation of genes involved upstream of the purine biosynthesis pathway (Figure S5), by which phosphoribosyl pyrophosphate (PRPP), the precursor for purine nucleotide synthesis produced from ribose-5-phosphate, is converted into inosine 5'-monophosphate (IMP). No significant concentration change was observed for metabolites in pyrimidine metabolism.

**Figure 2.**
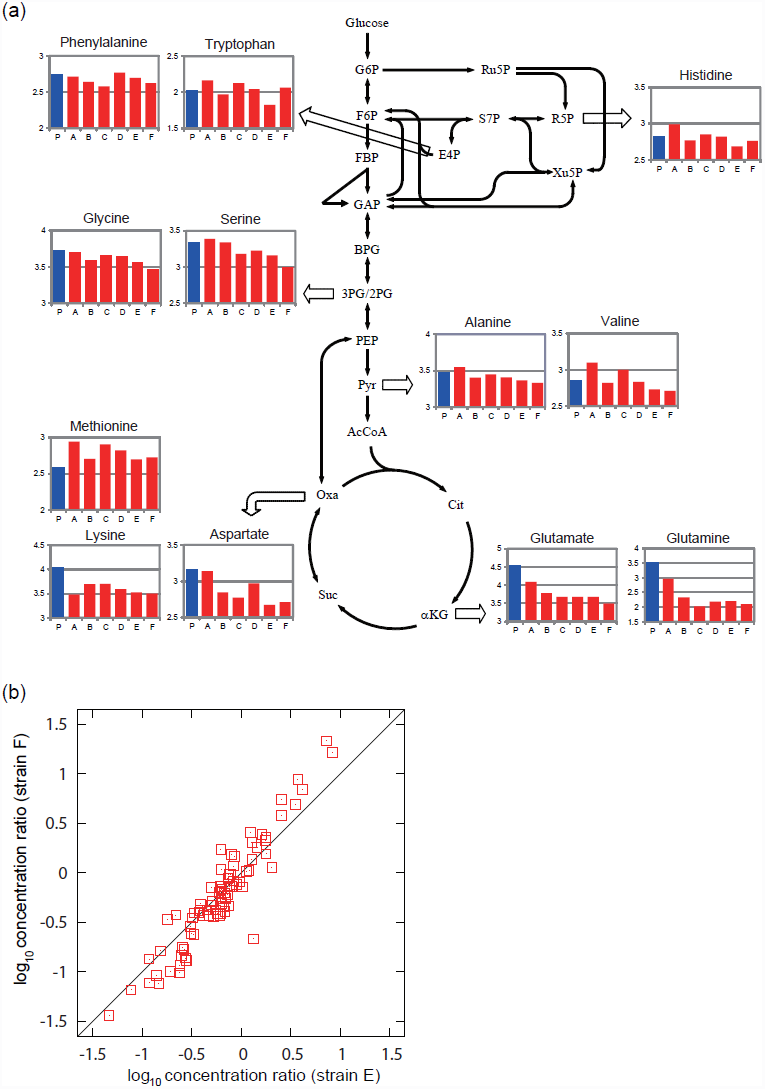
Metabolome analysis of ethanol tolerant *E. coli* strains. (a) Concentration of amino acids in ethanol tolerant strains. In each inset, the vertical axis shows the log-transformed absolute concentration (μM). (b) Correlation between metabolite concentration changes in strains E and F. Horizontal and vertical axes represent log-transformed concentration ratios with the parent strain, and each dot represents the concentration change of the metabolites.

The metabolome analysis also demonstrated similar changes in metabolite concentration among the tolerant strains, which resembles observations for gene expression changes. The correlation of overall metabolite concentration changes between independently obtained tolerant strains indicated similar metabolite shifts (Figure 2b and Figure S6). Both the transcriptome and metabolome analyses showed that phenotypic changes were similar among tolerant strains even though they were obtained from independent long-term cultivations.

### Genome resequencing analysis of ethanol tolerant strains

Genotype changes in each tolerant strain were analyzed using two high-throughput sequencers, SOLiD and Illumina Miseq (see Methods for details). In the resequencing analysis, we extracted genomic DNA samples from the cell population at the end-point of the experimental evolution without single-colony isolation, to identify genotype changes that were fixed in the majority of tolerant cells and to avoid a fixation of minority mutations. For point mutations, SOLiD and Miseq analyses identified 136 and 138 fixed mutations in all 6 tolerant strains, respectively, with 135 of these mutations being identified in both analyses. The discrepancy (4 point mutations in the strain A) was checked by Sanger sequencing, and it was confirmed that 3 were true positive and another was false positive. After screening indels by SOLiD sequencing, we identified 7 small (< 500 bp) and 13 large indels in all tolerant strains. We verified these small and large indels by Sanger sequencing, finding all were true positive. Finally, in strain C, the coverage of an approximately 200k-bp region was significantly higher than in other strains (Figure S7), which strongly suggested that the corresponding region duplicated during long-term cultivation.

The identified mutations at the end-point of experimental evolution are summarized in Table 1. The number of mutations in strain A was significantly larger than other strains (Table S3). The reason for the larger number of mutations was likely due to a mutation leading to a stop codon in the *mutS* coding region, which codes a mismatch repair protein. It is known that disruption of *mutS* significantly increases the mutation rate of *E. coli* cells (Glickman and Radman 1980). We confirmed that the number of mutations in strain A at 1512 h (about 600 generations) after commencing laboratory evolution was only three and that these did not include a mutation in *mutS*. This result suggested that after 1512 h, the mutation in the *mutS* gene was fixed and resulted in a significant increase in the mutation rate. The emergence of a strain with a significantly high mutation rate, or a 'mutator', is often observed in the laboratory evolution of microorganisms (Bachmann, et al. 2012; Barrick, et al. 2009; Levert, et al. 2010; Sniegowski, et al. 1997).

**Table 1:**

| Strain | Type | Gene | Position |  | Nucleotide | change | Gene Description |
| --- | --- | --- | --- | --- | --- | --- | --- |
|  |  |  | Reference | Gene |  |  |  |
| A | 125 SNPs and 6 Indels (see Table S3) |  |  |  |  |  |  |
| B | Ins | *hns* | 1294843 | -273 | 1195 bp | IS5 insertion, promoter | global DNA-binding transcriptional dual regulator |
|  | Ins | *yeaR* | 1881551 | 147 | – | IS186 insertion | conserved hypothetical protein |
|  | SNP | *iscR* | 2660496 | 292 | A→T |  | DNA-binding transcriptional activator |
|  | SNP | *ilvG* | 3685148 | 974 | A→T |  | acetolactate synthase II, large subunit |
| C | Del | 12 genes | 575013 |  | –6775 bp |  | *insH,nmpC,essD,ybcS,rzpD,rzoD,borD,ybcV,ybcW,nohB,tfaD,ybcY* |
|  | Ins | *nagE* | 705229 | 864 | +3:CCG | 3bp insertion | fused N-acetyl glucosamine specific PTS enzyme |
|  | Ins | *hns* | 1294843 | -273 | 1195 bp | IS5 insertion, promoter | global DNA-binding transcriptional dual regulator |
|  | SNP | *yeaY* | 1892079 | 168 | T→A | synonymous | predicted lipoprotein |
|  | Ins | *menC* | 2380504 | 485 | 1343 bp | IS186 insertion | o-succinylbenzoyl-CoA synthase |
|  | SNP | *relA* | 2910761 | 1547 | A→G |  | (p)ppGpp synthetase I/GTP pyrophosphokinase |
|  | SNP | *rpoC* | 3448513 | 2819 | G→A |  | RNA polymerase, beta prime subunit |
|  | SNP | *rpoA* | 4200347 | 961 | T→A |  | RNA polymerase, alpha subunit |
| D | Ins | *hns* | 1294843 | -273 | 1195 bp | IS5 insertion, promoter | global DNA-binding transcriptional dual regulator |
|  | SNP | *proQ* | 1916977 | 272 | A→T |  | predicted structural transport element |
|  | SNP | *ispG* | 2639469 | 992 | T→C |  | 1-hydroxy-2-methyl-2-(E)-butenyl 4-diphosphate synthase |
|  | SNP | *rpsD* | 4198966 | 226 | T→G |  | 30S ribosomal subunit protein S4 |
|  | Ins | *yjhA* | 4544220 | -39 | 1199 bp | IS5 insertion, promoter | N-acetylnuraminic acid outer membrane channel protein |
| E | Ins | *hns* | 1294843 | -273 | 1195 bp | IS5 insertion, promoter | global DNA-binding transcriptional dual regulator |
|  | Ins | *cspC* | 1909109 | 45 | 1199 bp | IS5 insertion | stress protein, member of the *CspA* (cold shock protein) family |
|  | SNP | *relA* | 2910944 | 1364 | A→T |  | (p)ppGpp synthetase I/GTP pyrophosphokinase |
|  | Ins | *yhcM* | 3379114 | 739 | +1:C |  | hypothetical protein with nucleoside triphosphate hydrolase domain |
|  | SNP | *atpE* | 3715545 | 54 | T→G |  | F0 sector of membrane-bound ATP synthase, subunit c |
| F | Del | *miaB* | 694563 | 728 | –88 bp |  | isopentenyl-adenosine A37 tRNA methylthiolase |
|  | Ins | *cspC* | 1909109 | 45 | 1199 bp | IS5 insertion | stress protein, member of the *CspA* (cold shock protein) family |
|  | SNP | *wzxC* | 2120992 | 1304 | A→T |  | colanic acid exporter |
|  | SNP | *iscR* | 2660468 | 320 | T→A |  | DNA-binding transcriptional activator |
|  | SNP | *relA* | 2911891 | 417 | T→G |  | (p)ppGpp synthetase I/GTP pyrophosphokinase |

In contrast to the more than hundred mutations fixed in strain A, the number of fixed mutations was relatively small in the other strains. As mentioned above, the phenotypic changes that occurred in independently evolved tolerant strains were similar, which might suggest mutations fixed in identical or related genes contributed to the changes. We found that mutations were commonly fixed in *relA*, which codes guanosine tetraphosphate synthetase. RelA regulates a stringent response by producing guanosine tetraphosphate (ppGpp) (Magnusson, et al. 2005). When *E. coli* cells face nutrient starvation or are exposed to various environmental stresses, the stringent response is signaled by ppGpp, which decreases growth-related activities including replication, transcription, and translation. Thus, the mutations commonly fixed in *relA* may relax the stringent response caused by ethanol stress to recover the growth activity. The mutations in *relA* and *spoT*, which codes an enzyme that plays a major role in ppGpp degradation, have been widely observed in the laboratory evolution of *E. coli* under various conditions, including glucose limitation (Cooper, et al. 2003) and high temperature (Kishimoto, et al. 2010). Here, relaxing the stringent response by mutating the *relA* and *spoT* genes may increase fitness under stress. Furthermore, in strains A, B, C, D, and E, insertion sequence element 5 (IS5) was inserted into the promoter region of *hns*, which codes for a DNA binding protein that has various effects on gene expression (Hulton, et al. 1990). However, no significant change of *hns* expression was observed in these strains. Except for *relA* and *hns*, no functional overlap among the mutations fixed in more than two tolerant strains was determined.

### Fitness contribution of fixed mutations

To evaluate the contribution of fixed mutations to the growth increase under ethanol stress, we introduced all identified mutations in strain F into the parent genome by site-directed mutagenesis (Posfai, et al. 1999). We identified 5 mutations, including 3 single nucleotide substitutions, one small deletion and one insertion in the genome of strain F. The sequence of the 1199-bp insertion was identical to IS5 and was inserted into and destroyed the *cspC* gene of strain F. Since the insertion of IS5 into the same position of the parent genome was difficult experimentally, we constructed a *cspC* deletion strain. Figure 3 shows the growth rates of the constructed strains by site-directed mutagenesis measured under the ethanol stress condition. The results demonstrated that the fixed mutation in *relA* significantly contributed to the growth rate increase (p<0.05; determined by t-test). However, other mutations had no significant effect on the growth rate, and even when all fixed mutations in strain F were introduced into the parent genome, the observed growth rate of the mutated strain was significantly smaller than that of strain F under the ethanol stress condition. These results suggested that the growth increase observed in strain F cannot be fully explained by the fixed mutations.

**Figure 3.**
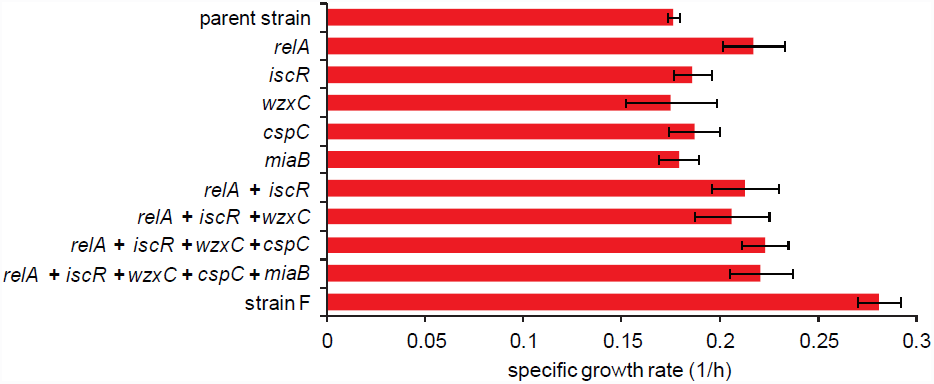
Growth rates of site-directed mutants. All mutations identified in strain F were introduced back to the parent strain. For each mutant, the names of the mutated genes are shown. Error bars indicate standard deviations calculated from three independent cultures.

### Timing of fixed mutations

To further evaluate the contribution of the fixed mutations in strain F on ethanol tolerance, we analyzed the relationship between the growth increase under ethanol stress and the timing of mutation-fixation events in long-term cultivation. To identify the timing, genomic DNA samples obtained at 12 different time points were applied to Sanger sequencing. The genomic DNA samples were obtained from cell populations that had heterogeneous genotypes. Thus, in some cases, the peak signals in the Sanger sequencing revealed mixed populations, i.e., cells with and without a specific mutation coexisted in the population. Figure 4a shows the time the mutations in strain F emerged. The increased growth rate did not always correlate with fixation events. More importantly, although at 576 h after inoculation no mutation was fixed in the majority of the cell population, the growth rate under ethanol stress significantly increased. Some cells at 576 h had mutations in the *relA* and *cspC* genes that may have contributed to the observed growth increase. To confirm this possibility, we isolated 48 clones from the cell population at 576 h and analyzed fixed mutations in *relA* and *cspC* by Sanger sequencing. Among the 48 clones, 5 had both *relA* and *cspC* mutations, 6 had the *cspC* mutation only, and the other 37 clones had no mutation. To evaluate the effect of these mutations on the population at 576 h, we measured the growth rates of clones with and without mutations under the ethanol stress condition. Clones with or without *relA* and *cspC* mutations showed significantly larger growth rates than parent strains (p<0.05, t-test), and there was no significant growth difference between clones (Figure 4b). These results indicated that the growth increase from 216 to 576 h cannot be explained by mutations in *relA* or *cspC*.

**Figure 4.**
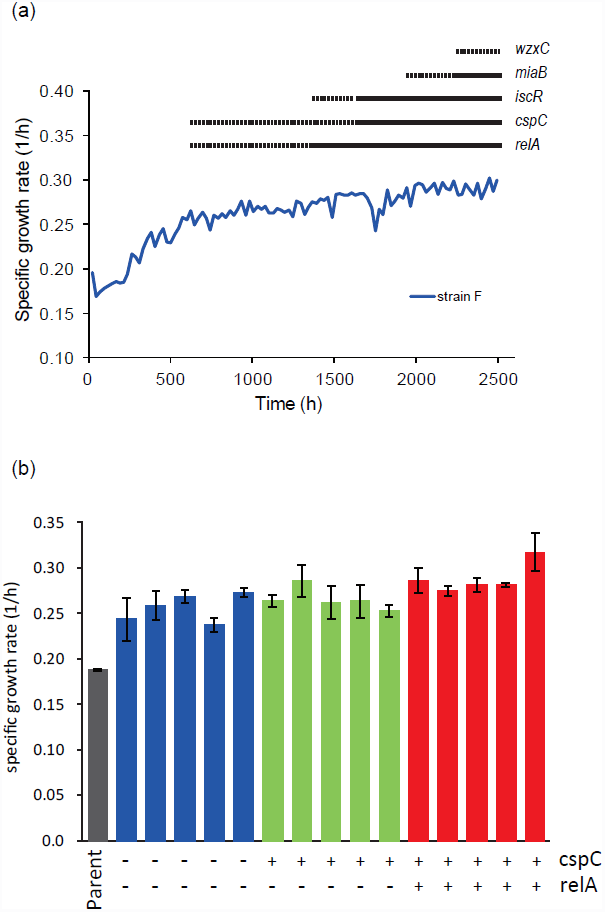
Time-series analysis of the mutation fixation. (a) Timing of mutation fixation events in strain F. To identify the timing of mutation fixation, genomic DNA samples obtained at 12 different time points (216, 384, 576, 744, 888, 1056, 1224, 1392, 1584, 1824, 1968, and 2232 hours after inoculation) were applied to Sanger sequencing. For each of the 5 identified mutations, the results of the Sanger sequencing is presented as a solid or dotted line. The solid line indicates that the mutation was fixed in the population at the corresponding time point, while the dashed line indicates the case of two peak signals, which indicates a mixed population of cells with and without the mutation. For example, in cells at 576 hours after inoculation, only some have mutations in *relA* and *cspC.* (b) Specific growth rates of cloned *E. coli* cells with and without *cspC* and *relA* mutations. The clones were isolated from strain F cell populations at 576 h. Each bar represents the specific growth rate of an isolated clone, where "+" and "-" mean with and without the corresponding mutation, respectively. Blue, green, and red bars represent the growth rates of clones without mutations, that with *cspC* mutation only, and with *cspC* and *relA* mutations, respectively. Error bars indicate standard deviations calculated from three independent cultures.

### Stability of ethanol tolerance

If the observed ethanol tolerance was due to phenotypic plasticity without genetic alteration, the phenotype of the ethanol tolerance would likely be unstable when the environment changes. We therefore cultivated cells with ethanol tolerance in an ethanol-stress free environment for 100 generations. Two cell populations were used: strain F and the cell population obtained at 576 h in the cultivation of strain F. After cultivation in the non-stress condition, we measured the growth rate under 5% ethanol stress to evaluate the stability of the ethanol tolerance. Ethanol tolerance was stably maintained even after 100 generations (Figure S8), which suggests that the observed phenotypic changes in the tolerant strains were stably memorized and passed to progeny cells.

### Discussion

Transcriptome and metabolome analyses revealed that phenotypic changes that occurred in ethanol tolerant strains were similar among independently evolved strains. Gene expression changes over time were found to exhibit high similarity among tolerant strains, which included non-monotonic expression changes with time scales much longer than the generation time. Such synchronized slow expression changes are difficult to explain by phenotypic changes caused by a small number of mutations. Instead, the observed phenotypic convergence to similar orbits more likely suggests the existence of deterministic slow dynamics that acquire ethanol tolerance.

Using high-throughput sequencers, we identified fixed mutations in the tolerant strains. One tolerant strain had a significantly larger number of fixed mutations than the others, probably due to disruption of *mutS*, which is involved in the mismatch repair mechanism. For the other tolerant strains, the number of fixed mutations was less than 10. We found that these mutations were commonly observed in the *relA* gene, which is involved in stringent response via ppGpp production, suggesting that the stringent response triggered by the ethanol stress was relaxed by these mutations in the tolerant strains and therefore did not diminish growth activity as would otherwise be expected. These mutations could be regarded as candidate beneficial mutations for ethanol tolerance. However, their small number suggests they are unlikely to explain the synchronized slow expression changes mentioned above. Consistent with this conclusion, we introduced these mutations into the genome of the parent strain of strain F and confirmed that the observed ethanol tolerance could not be reproduced (Figure 3). This result suggested that phenotypic plasticity without genetic alteration contributed to the observed ethanol tolerance.

That phenotypic plasticity contributed to ethanol tolerance was also supported by the timing of the mutation fixation. The increase in growth rate of strain F did not correlate with mutation fixation events, and *E. coli* clones without any beneficial mutation grew significantly faster than the parent strain under ethanol stress (Figure 4). Importantly, the phenotype of ethanol tolerance in these strains was stably maintained even when the environmental conditions changed, and the ethanol tolerance was maintained after cultivation of 100 generations under a condition without ethanol stress.

Based on these results, we propose that part of the growth increase observed in the adaptive evolution experiment under ethanol stress was due to phenotypic plasticity without genomic alteration, and that this plasticity could be stably memorized in the intra-cellular state and be inherited by progeny cells. At present, the mechanism for this non-genetic memory of adaptive phenotypic change is unknown. Previous studies have demonstrated that fluctuations in the expression dynamics can contribute to the acquisition of antibiotic resistance (Wakamoto, et al. 2013). However, in such cases, an adapted state is generally maintained only for several generations, since information on the adapted state decays with each cell division. In contrast, that ethanol tolerance was maintained in tens of generations in the present study suggests machinery for information inheritance. Similar epigenetic memory was also suggested to play a role in the evolution of antibiotic resistance in *E. coli* (Adam, et al. 2008). In *E. coli* cells, genome methylation patterns are known to act as epigenetic memory that controls the expression profile (Heithoff, et al. 1999; Palmer and Marinus 1994), as too is the binding of histone-like proteins, such as H-NS and Fis, to genomic DNA (Gonzalez-Gil, et al. 1996; Williams and Rimsky 1997). These epigenetic mechanisms might contribute to the observed non-genetic memory and should be considered in future works.

In conclusion, we analyzed phenotypic and genotypic changes of *E. coli* cells that occurred during adaptive evolution to ethanol stress and found that at least part of the adaptive phenotypic changes were not due to genomic mutations. The phenotype of ethanol tolerance was stably maintained after environmental changes, which might suggest that a mechanism for non-genetic memory plays a role in the adaptation process. Of course, it is difficult to completely exclude the possibility that genomic mutations can explain the acquisition of observed tolerance. For example, mutations disappeared in the end-point population might contribute to the fitness under ethanol stress. Thus, complete understanding of the mechanisms responsible for adaptive evolution will require detailed study of both genetic and non-genetic contributions to adaptive phenotypic changes and their interplay.

## Materials and Methods

### Laboratory evolution

The *E coli* strain W3110 was obtained from National BioResource Project (National Institute of Genetics, Japan) and used for all laboratory evolution cultures. Ethanol tolerant strains, A through F, were obtained as previously described (Horinouchi, et al. 2010). Briefly, cells were grown in 10 ml of M9 minimal medium with 5% (v/v) ethanol at final concentration. Cell cultures were performed at 30 °C with shaking at 150 strokes min^−1^ using water bath shakers. We diluted the cells in fresh medium every 24 hours and maintained an exponential growth phase by adjusting the initial cell concentration.

### Transcriptome analysis by microarray

For transcriptome analysis, a custom-designed tilling microarray of *E. coli* W3110 in Affymetrix platform was used. The platform contained approximately 1.5 million perfect-match 21-bp probes for the *E. coli* genome and an approximately 4.5 million of corresponding single-base mismatch probes (Ono, et al. 2013). For the sample preparation, each strain was inoculated from the frozen stock into 10 mL of M9 medium for preculture. Five-microliter aliquots of preculture medium cells were inoculated into 10 mL of M9 medium with or without 5% (v/v) ethanol and cultured for 10 generations (with ethanol) or 5 generations (without ethanol). Cells in the exponential growth phase were harvested by centrifugation and stored at −80 °C before RNA extraction. Total RNA was isolated and purified from cells using an RNeasy mini kit with on-column DNA digestion (Qiagen, Hilden, Germany). The synthesis of cDNA, fragmentation and end-terminus biotin labeling were carried out in accordance with Affymetrix protocols. Hybridization, washing, staining, and scanning were carried out according to the Expression Analysis Technical Manual (provided by Affymetrix). To obtain the absolute expression levels of genes from microarray raw data, we used the Finite Hybridization model (Furusawa, et al. 2009; Ono, et al. 2008). Expression levels were normalized using the quantile normalization method (Bolstad et al., 2003). Information on gene regulation was obtained from RegulonDB (Gama-Castro et al., 2008). Both the normalized expression data sets and the raw CEL files were deposited in the NCBI Gene Expression Omnibus database under the GEO Series accession number GSE59050.

### Metabolome analysis by capillary electrophoresis time-of-flight mass spectrometry

Metabolomic analysis was performed using capillary electrophoresis time-of-flight mass spectrometry (CE-TOFMS). The sample preparation method for CE-TOFMS analysis was previously reported (Yoshikawa, et al. 2013). Briefly, cells in the exponential growth phase were harvested by filtration (Isopore™ Membrane Filters HTTP, Millipore, Billerica, MA) and washed with water. The filter was immersed in methanol containing internal standards to quench metabolic reactions and extract intracellular metabolites before sonication for 30 s. To remove phospholipids, the methanol solution was mixed with chloroform and water and then centrifuged at 4,600 g for 5 min at 4 °C. The separated methanol/water layer was filtered through a 5 kDa cutoff filter (Millipore) by centrifugation at 9,100 g and 4 °C to remove proteins. The filtrate was lyophilized and dissolved in 25 μL of water prior to the CE-TOFMS analysis.

CE-TOFMS analysis was performed using the Agilent 7100 CE system equipped with the Agilent 6224 TOF-MS system, the Agilent 1200 isocratic HPLC pump, the G1603A Agilent CE/MS adapter kit, and the G1607A Agilent CE/MS sprayer kit (Agilent Technologies). For system control and data acquisition, Chemstation software for CETOFMS (Agilent Technologies) and MassHunter software (Agilent Technologies) were used. The concentration of each metabolite in methanol was quantified using the relative peak area of each metabolite to the internal standard peak area obtained from biological samples and the relative peak area obtained from chemical standards mixtures that included amino acids; intermediate metabolites from glycolysis, TCA cycle, and PPP (50 μM each); and internal standards including 25 μM methionine sulfone and 25 μM camphor-10-sulfonic acid (Human Metabolome Technologies) analyzed in parallel with experimental samples. Peak area data were obtained using the MassHunter software for qualitative analysis (Agilent Technologies).

### Genome resequencing

Frozen stocks of the strains were grown overnight in 10 ml of M9 minimal medium at 30 °C. Precultured cells were diluted to OD600nm 0.05 and grown in 10 mL of fresh M9 medium. When OD600 nm reached approximately 2.0, Rifampicin (final concentration 300 μg/mL) was added to block the initiation of DNA replication, and the culture was continued for another 3 hours. The cells were collected by centrifugation at 16,000 × g for 2 min and 25 °C and then the pelleted cells were stored at −80 °C prior to genomic DNA purification. Genomic DNA was isolated and purified using a Wizard^®^ Genomic DNA Purification kit (Promega) in accordance with the manufacturer's instructions. To improve the purity of genomic DNA, additional phenol extractions were performed before and after the RNase treatment step. The purified genomic DNAs were stored at −30 °C prior to use.

The same genomic DNA samples of the parent and ethanol tolerant strains were sequenced using both SOLiD DNA analyzer (Life Technologies) and Illumina MiSeq Desktop Sequencer (Illumina). For SOLiD sequencing, mate-paired libraries (2x50 bp) of 1200 bp insert size were generated and sequenced according to the manufacturer's protocol, which resulted in about 200-fold coverage on average. For Illumina sequencing, paired-end libraries (2x250 bp) were generated using Nextera v2 technology and sequenced by the MiSeq system according to the manufacturer's protocol (Illumina), which resulted in about 180-fold coverage on average. The mate-pair sequencing data by SOLiD and the paired-end sequencing data by Illumina Miseq are available from the DDBJ Sequence Read Archive of the DNA Data Bank of Japan (DRA) under accession number DRA002309.

For identification of point mutations by SOLiD sequencing, the sequence reads were mapped to the reference genome of *E. coli* W3110 with SOLiD bioscope software (version 1.2.1) (Life Technologies). Point mutations were subsequently called by the diBayes algorithm (Life Technologies), in which the threshold p-value was set to 10^−7^. To obtain only those mutations present in the majority of cells, variant calls with a ratio of variant reads less than 0.6 were excluded from further analysis. For Illumina sequence data, the sequence reads were mapped to the reference genome by SSAHA2 (Ning, et al. 2001). For each potential point mutation, we extracted those with coverage reads more than 10 and a ratio of variant read to wild-type read more than 0.6. When the point mutation calls by these two methods produced discrepancies, the candidate mutations were confirmed by Sanger sequencing.

The identification of small indels (< 500 bp) were performed by SOLiD bioscope software, in which the default parameter setting was used. The small indels identified by SOLiD sequencing and bioscope software were confirmed visually using the mapping of reads obtained by Illumina MiSeq.

For the identification of large indels, we implemented a detection algorithm based on distances between mapped SOLiD mate-paired sequence reads as follows. After removing low quality reads (mapping quality < 10 or including bases with base quality < 30), we mapped mate-paired sequence reads by bioscope software, and then used all mapped read pairs to calculate the mean and standard deviation of the distance between any two mapped reads. When indels are fixed in the genome, the distance between two mapped reads mapped to one region shows a deviation from the other genome region. We screened genomic regions at which the median of the mapped read distances was more than 3 SD from the mean, and the presence of an indel was confirmed visually using the mapping. When the pattern of the read distances suggested an insertion and part of the counterpart reads was mapped to an IS element, an IS element insertion was assumed and validated manually. All indels identified by SOLiD sequencing were also identified by Sanger sequencing, as predicted.

The sequence reads from the parent strains were also mapped to the reference genome of *E. coli* W3110, and mutations were screened by the above methods. Point mutations and indels found in the parent strains were also found in all tolerant strains and discarded from further analysis.

### Effect of genomic mutations on ethanol tolerance

Each identified mutation was introduced into the parent strain using the suicide plasmid method (Posfai, et al. 1999). This approach enables the introduction of any desired mutation without leaving an antibiotic marker in the genome. DNA fragments including identified mutations were cloned into suicide plasmid pST76-K and inserted into the chromosome of the parent strain. Allele replacement and marker removal was performed using helper plasmid pUC19RP12 (These plasmids were kind gifts from Dr. Gyorgy Posfai, Biological Research Centre of the Hungarian Academy of Sciences, Hungary). To eliminate the helper plasmid, obtained mutants were cultured in M9 medium at 30 °C. Primer information of the mutant construction is summarized in Supplementary Table S4. To evaluate the effect of the mutations on ethanol tolerance, the specific growth rate of mutated strains was quantified in M9 medium with 5% ethanol. The conditions for these cultures were identical to those in laboratory evolution. The cultures of each strain were performed three times independently.

## Acknowledgements

We thank Dr. Peter Karagiannis for proofreading of the manuscript, and Ms. Yuki Hidaka, Kumi Takabe for technical assistance. This work was supported in part by Grant-in-Aid for Young Scientists (A) [23680030 to C.F.], Grant-in-Aid for Scientific Research (A) [24246134 to H.S.], and Grant-in-Aid for Young Scientists (B) [24700305 to SS] from JSPS, and Grant-in-Aid for Scientific Research on Innovative Areas [25128715 and 23128509 to C.F.] from MEXT, Japan.

## Author Contributions

THo, TY, HS, and CF conceived and designed the experiments. THo, SS, and THi performed the experiments. THo, NO, and CF analyzed the data. THo and CF wrote the manuscript. All authors contributed to data interpretation, reviewed the manuscript, and approved the final version.

---

**Figure S1.**
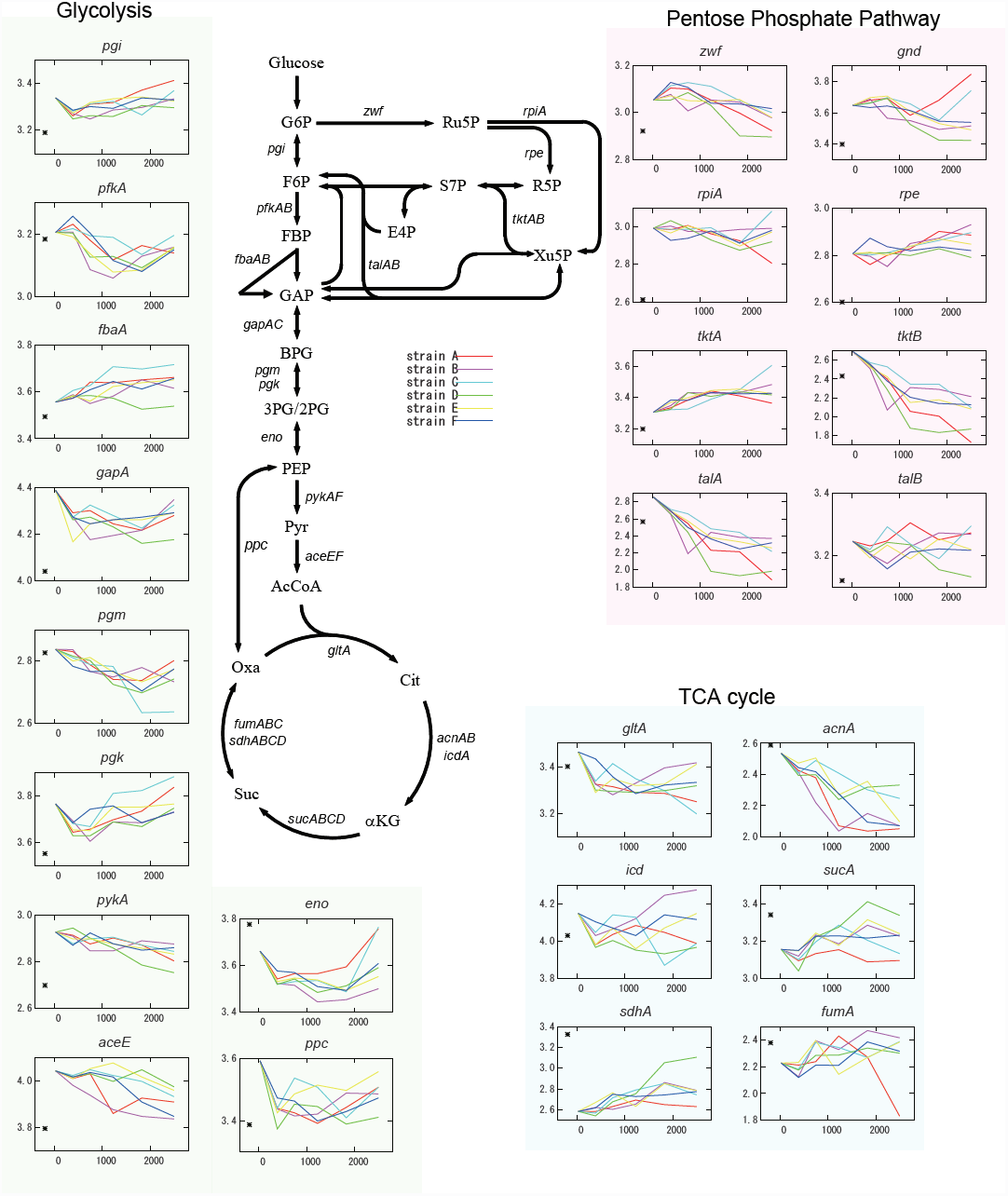
Expression changes of genes related to glycolysis, the pentose phosphate pathway, and TCA cycle. The horizontal axis shows time (hours), and the vertical axis shows expression level (a.u.). Asterisks (*) in the insets indicate expression levels obtained in the absence of ethanol stress for reference. Abbreviations: 2PG, 2-Phosphoglyceric acid; 3PG, 3-phosphoglycerate; AcCoA, acetyl-CoA; α KG, α -ketoglutarate; BPG, 1,3-bisphosphoglycerate; Cit, citrate; E4P, erythrose4-phosphate; F6P, fructose 6-phosphate; FBP, fructose 1,6-bisphosphate; GAP, glyceraldehyde 3-phosphate; G6P, glucose 6-phosphate; Oxa, oxaloacetate; PEP, phosphoenolpyruvate; Pyr, pyruvate; R5P, ribose 5-phosphate; S7P, sedoheptulose 7-phosphate; Suc, succinate; X5P, xylulose 5-phosphate.

**Figure S2.**
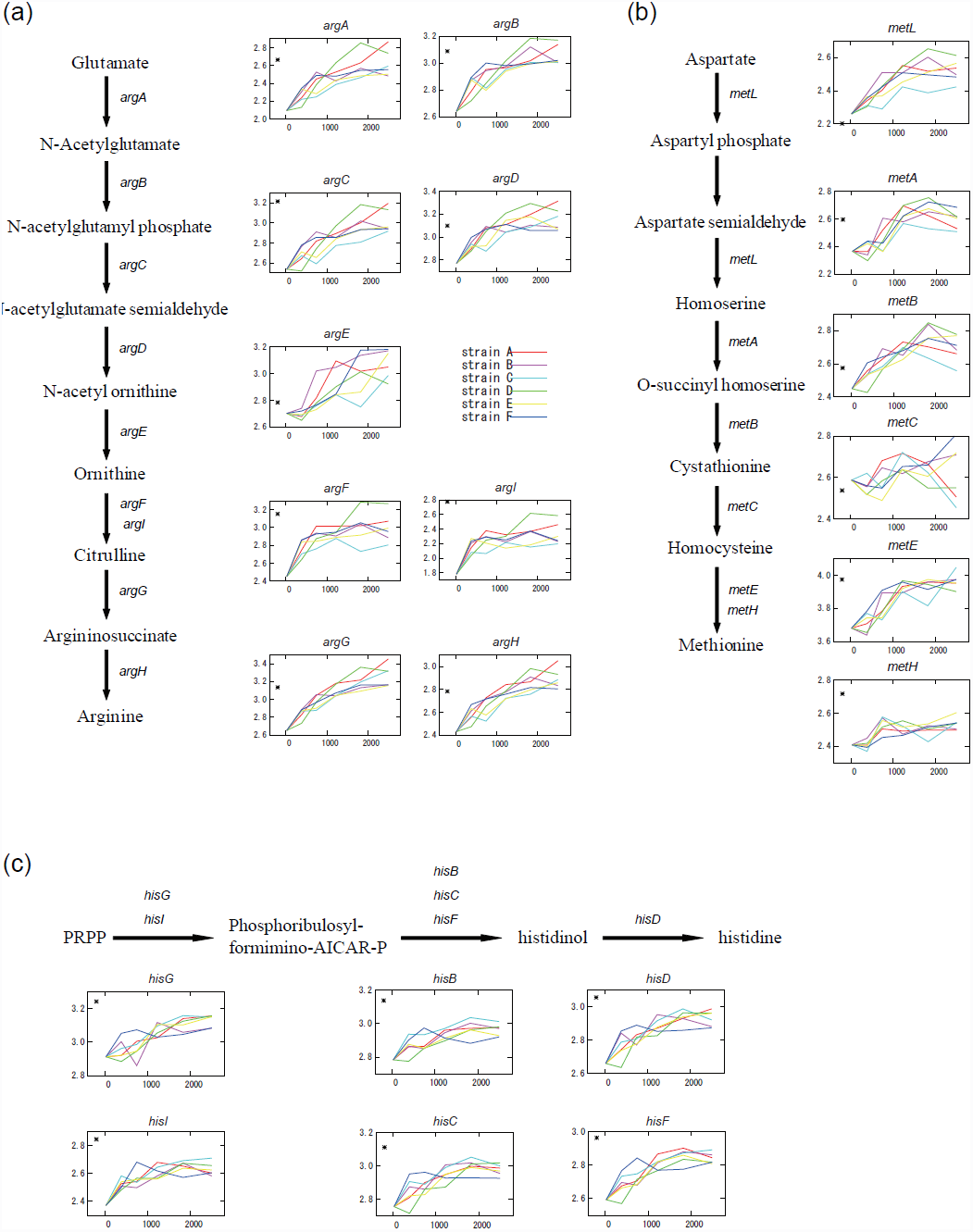
Expression changes of genes related to (a) arginine, (b) methionine, and (c) histidine biosynthesis pathways in tolerant strains. Abbreviations: PRPP, phosphoribosyl pyrophosphate.

**Figure S3.**
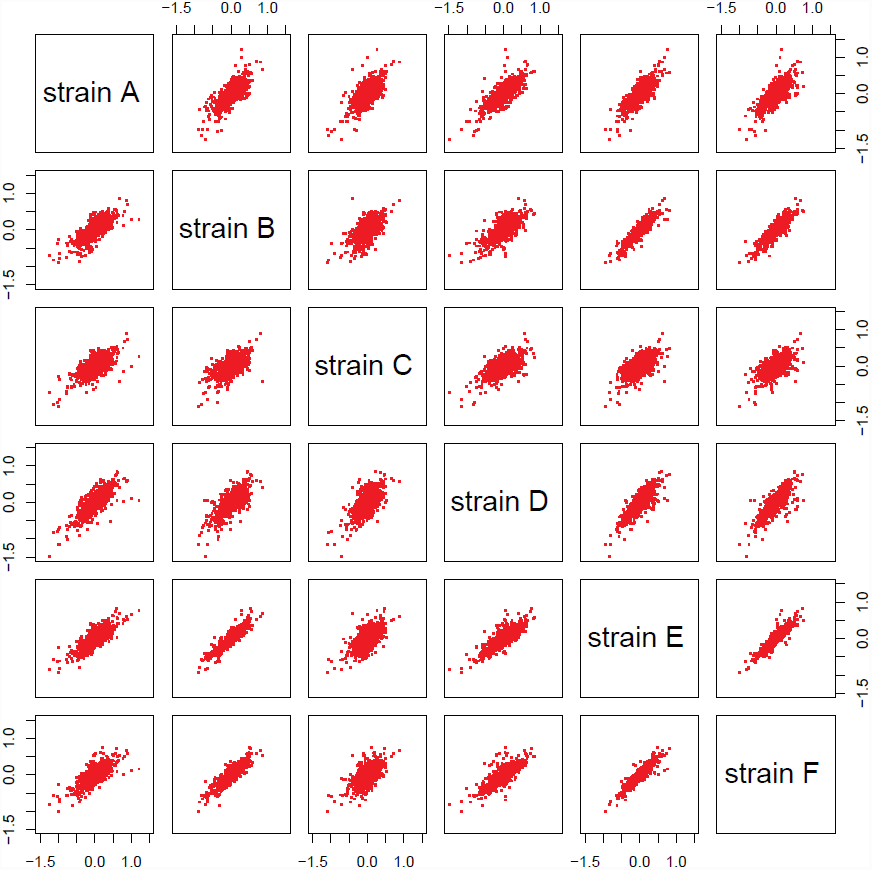
Correlations between gene expression changes for all possible pairs of tolerant strains. Each axis represents log_10_-transformed expression changes between a tolerant strain and the corresponding parent strain, while each dot represents the expression changes of a gene.

**Figure S4.**
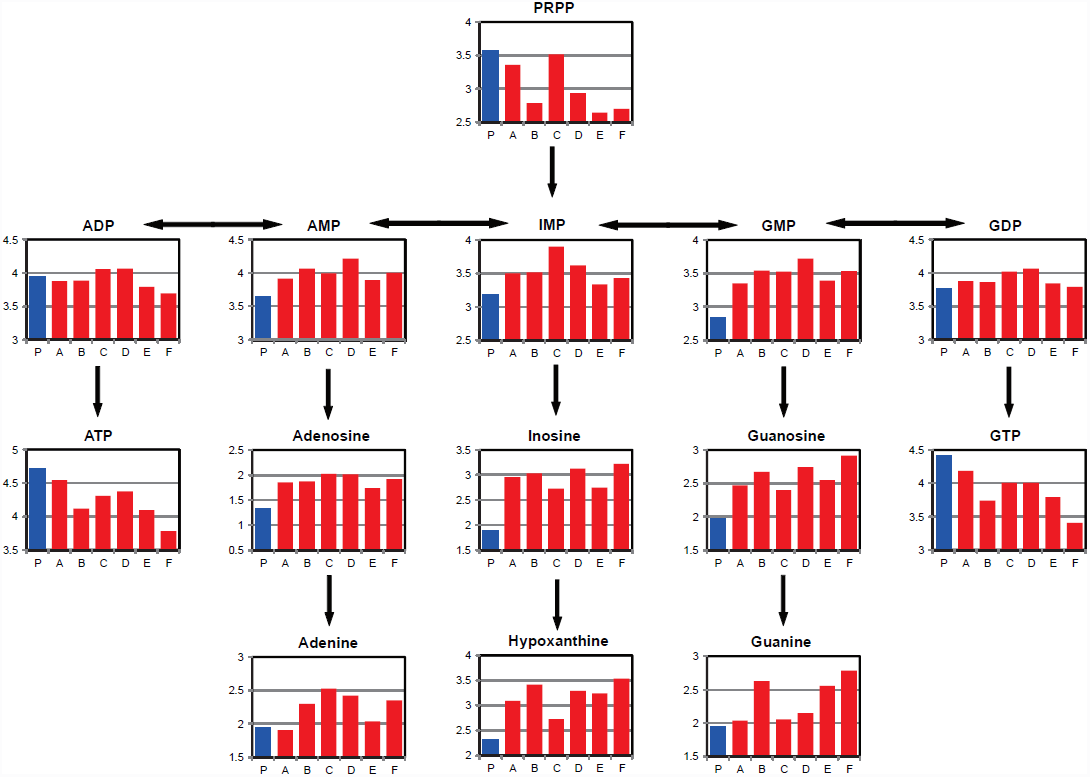
Metabolite concentrations in *de novo* and salvage purine biosynthesis. In each inset, the vertical axis shows the log-transformed absolute concentration (μM). Abbreviations: AMP, adenosine monophosphate; ADP, adenosine diphosphate; ATP, adenosine triphosphate; GMP, guanosine monophosphate; GDP, guanosine diphosphate; GTP, guanosine triphosphate; IMP, inosine monophosphate.

**Figure S5.**
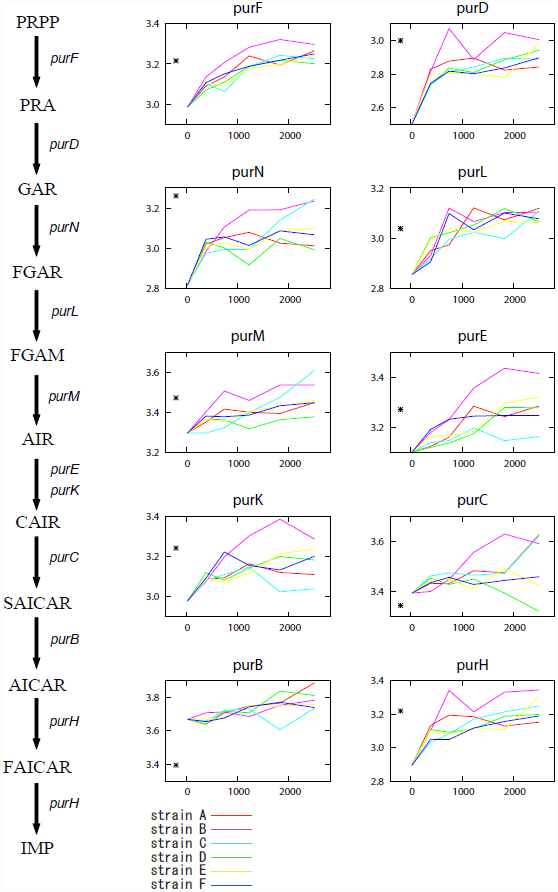
Expression changes of genes related to the biosynthesis of phosphoribosyl pyrophosphate (PRPP) in tolerant strains. Abbreviations: PRA, 5-phospho-β-D-ribosylamine; GAR, N^1^-(5-phospho-β-D-ribosyl)glycinamide; FAGR, N^2^-formyl-N^1^-(5-phospho-β-D-ribosyl)glycinamide; FAGM, 2-(formamido)-N^1^-(5-phospho-β-D-ribosyl)acetamidine; AIR, 5-amino-1-(5-phospho-D-ribosyl)imidazole; CAIR, 5-amino-1-(5-phospho-D-ribosyl)imidazole-4-carboxylate; SAICAR, (S)-2-[5-amino-1-(5-phospho-D-ribosyl)imidazole-4-carboxamido]succinate; AICAR, 5-amino-1-(5-phospho-D-ribosyl)imidazole-4-carboxamide; FAICAR, 5-formamido-1-(5-phospho-D-ribosyl)imidazole-4-carboxamide.

**Figure S6.**
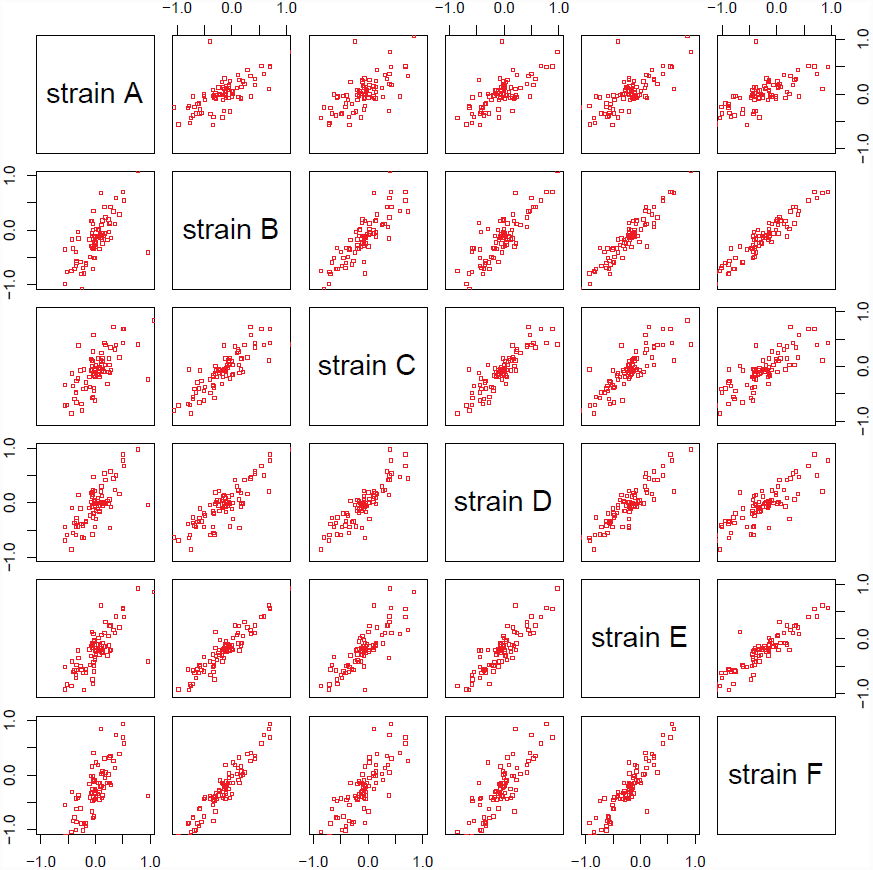
Correlations between metabolite concentration changes for all possible pairs of tolerant strains. Each axis represents log_10_-transformed metabolite concentration changes between a tolerant strain and corresponding parent strain, while each dot represents the concentration changes of a metabolite.

**Figure S7.**
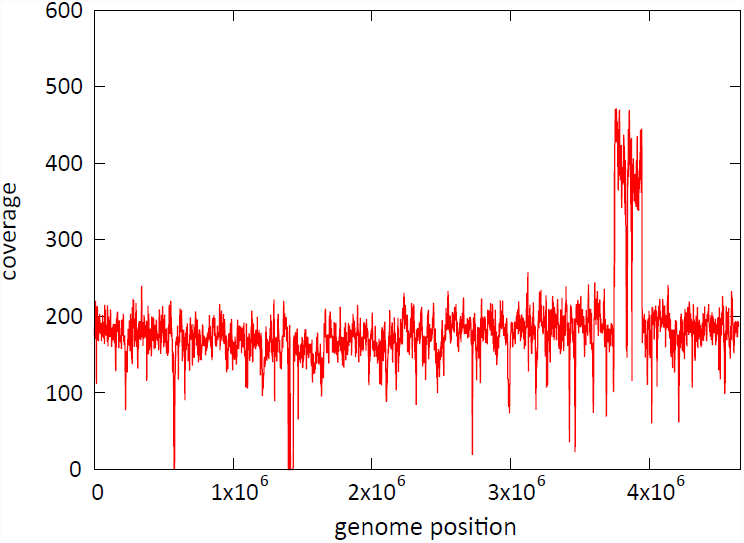
Sequence coverage of strain C. The number of mapped sequencing reads of Illumina analysis is plotted as a function of genome position. The coverage almost doubled in the region from 3750000 to 3950000 bp in the W3110 reference genome position, suggesting genomics duplication. The region includes 186 genes. No similar duplication was observed in other tolerant strains.

**Figure S8.**
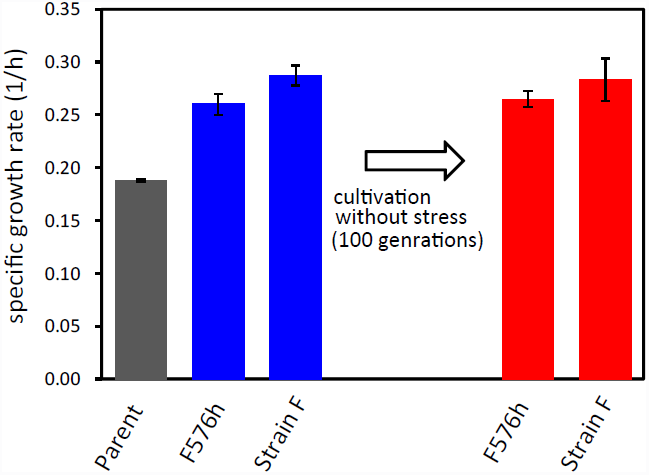
Stability of ethanol tolerance. Strain F at the end point (2500 h) and at 576 h was cultivated for 200 generations absent ethanol stress. After the cultivation, ethanol tolerance was evaluated by measuring specific growth rates in 5% ethanol stress (red bars). The growth rates under ethanol stress were similar to those before the non-stress cultivation (blue bars) and were significantly higher than that of the parent strain.

